# Abundance-weighted analysis and deep functional metagenomic selection reveals unrecognized diversity of soil-derived streptothricin acetyltransferases

**DOI:** 10.64898/2026.10.29.762622

**Authors:** Lufan Yang, Terence S. Crofts

**Affiliations:** Department of Animal Science, University of Illinois Urbana-Champaign, Urbana, IL, United States; Carl R. Woese Institute for Genomic Biology, University of Illinois Urbana-Champaign, Urbana, IL; Division of Nutritional Sciences, University of Illinois Urbana-Champaign, Urbana, IL

## Abstract

Streptothricins (STCs) are broad-spectrum antibiotics under investigation for their therapeutic potential against antibiotic-resistant Gram-negative pathogens. However, the extent and diversity of genes that confer resistance to STCs, or their environmental resistome, remains largely unknown and is essentially limited to a handful of STC acetyltransferases. Here, we used a deep functional metagenomic approach to uncover previously unrecognized STC resistance determinants. A functional metagenomic library containing *ca.* 700 Gigabases of soil metagenomic DNA was selected for resistance to nourseothricin (NTC), a mixture of STC congeners, resulting in collection and sequencing of nearly 30,000 resistant clones. An abundance-weighted sequence analysis identified 265 acetyltransferase-like candidates spanning broad phylogenetic, taxonomic, and genomic diversity. Re-cloning and experimental testing of 18 candidates chosen to cover acetyltransferase sequence space confirmed that 10 conferred significant NTC resistance in *Escherichia coli*. Phylogenetic analyses identified three previously uncharacterized STC acetyltransferase lineages, designated as SatC, SatD, and SatE. Despite substantial sequence divergence from characterized streptothricin acetyltransferases (maximum amino acid identities of approximately 31% for SatC and SatD and up to 47.5% for SatE), these enzymes largely retained key residues associated with streptothricin binding and catalysis. In addition to strongly increasing NTC resistance, expression of the SatC construct in *E. coli* also significantly decreased susceptibility to the aminoglycoside antibiotics amikacin and kanamycin. Together, these findings reveal a substantially broader than previously known environmental repertoire of STC acetyltransferases and demonstrate how deep functional metagenomic selections, combined with abundance-weighted hit calling, phylogenetic analysis, and structural prediction, can uncover novel resistance functions.

## INTRODUCTION

Antimicrobial resistance (AMR) challenges the effectiveness of existing clinical antibiotics while the development pipeline of new antibacterials remains slow(1, 2). In 2019, bacterial AMR was associated with an estimated 4.95 million deaths worldwide, including 1.27 million deaths directly attributable to resistance(2). The majority of antibiotics are natural products or their derivatives(3, 4) and it is therefore unsurprising that antibiotic resistance is a natural and ancient property of microbial communities(5, 6). The complete collection of resistance genes in a given microbiome, including pathogenic, commensal, and environmental organisms, is its resistome. The antibiotic resistome includes *bona fide* antibiotic resistance genes as well as genes that may evolve into resistance genes or gain the ability to confer resistance if their genomic context changes (6). Natural products therefore remain important sources for the discovery of new antibiotics but research into the mechanisms of drug resistance that co-occur with them in nature is a matter of urgency.

The streptothricin antibiotics (STCs) are natural product antibiotics which were discovered more than 80 years ago(7, 8). Streptothricin was first isolated in 1942 from a soil-derived *Streptomyces lavendulae* strain and was recognized for activity against both Gram-positive and Gram-negative bacteria(9). STCs share a streptolidine lactam linked to a carbamoylated gulosamine sugar and differ from each other in the length of their β-lysine homopolymer chains. These chains range from a single residue in streptothricin F (STC-F) to longer chains in other congeners (STC-A through STC-E and STC-X)(7, 10) (**Fig. 1A**). Although STCs share a bactericidal translational miscoding mechanism of action with the aminoglycoside family of antibiotics they are chemically distinct and target a different region of the ribosome (16S rRNA helix 34 for streptothricins and helix 44 for many aminoglycosides) and should be classified as their own antimicrobial class (7, 11, 12). Despite their potential clinical utility, STC antibiotics have only been used in agriculture due to observed nephrotoxicity in humans and other mammals (7).

**Figure 1.**
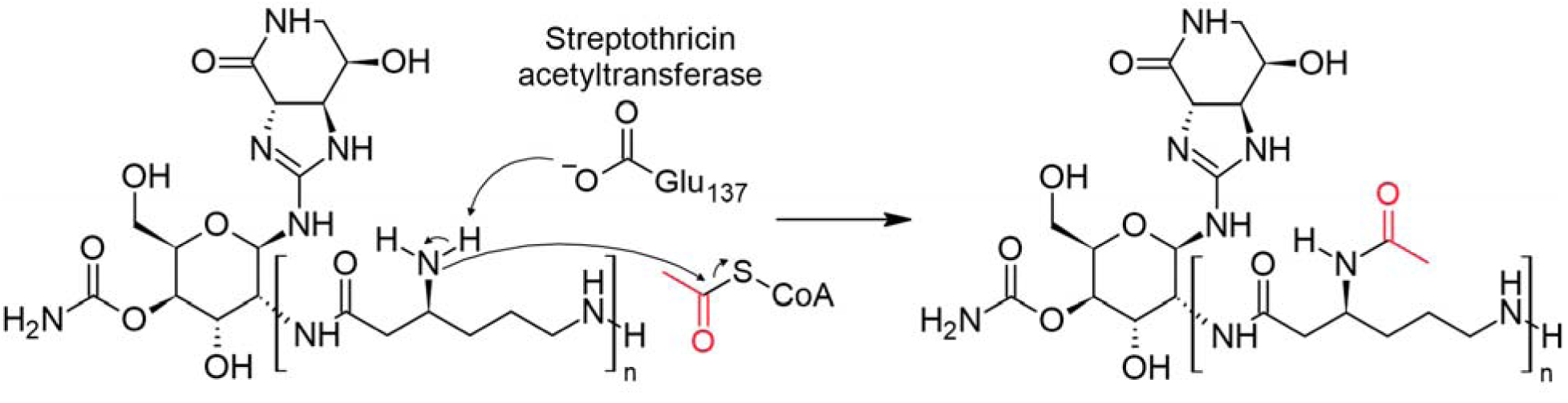
Chemical structure of streptothricin antibiotics and mechanism of acetyltransferase mediated resistance. Streptothricin antibiotics consist of a streptolidine lactam ring, a gulosamine sugar, and a variable length β-lysine chain (indicated by brackets), with the number of β-lysine residues determining the streptothricin variant. Streptothricin acetyltransferases (SATs) catalyze the transfer of an acetyl group from acetyl-CoA to the β-amino group of the β-lysine moiety. A conserved glutamate residue (Glu137 in the enzyme shown) acts as a general base, facilitating deprotonation of the β-lysine amino group during acetyl transfer. The transferred acetyl group is highlighted in red.

Recently, Morgan *et. al.,* determined that STC-F has lower toxicity than longer-chain STCs, while still being effectively bactericidal against multidrug-resistant Gram-negative bacteria (12). This potential for an improved therapeutic window and the recent publication of a total synthesis of STC-F (making synthetic modifications more feasible) has increased interest in the clinical potential of this antibiotic class (7, 10). STC production is widespread among soil *actinomycetes*, with phenotypic screens frequently finding that ∼40% of tested actinomycete strains are capable of producing STC-like substances (7). However, while soil microbial communities are major sources of natural-product antibiotics, they are also diverse reservoirs of antibiotic resistance (13, 14). Despite the abundance of STC-producing bacteria in soil, only a few STC resistance genes have been experimentally characterized to date. This discrepancy suggests the existence of a largely unexplored environmental streptothricin resistome, with known resistance genes representing only a fraction of its functional diversity (7). As progress is made towards bringing STC antibiotics into the clinic, understanding the potential pool of resistance mechanisms that might combat them is important.

Inactivation by acetylation is the main characterized mechanism of STC resistance, with the resistance-associated modification occurring at the β-amino group of the β-lysine moiety(15)(**Fig. 1**). STC acetyltransferases including STAT, SAT-2, SAT-3, SAT-4, SatA, SatB, SrStat1, and SrStat2(16–23) have been shown to inactivate STC and confer resistance. All these enzymes belong to the large Gcn5-related N-acetyltransferase (GNAT) superfamily, whose members share a conserved structural core and cofactor-binding architecture. Despite this shared structure, substrate recognition can vary substantially among GNAT enzymes, and proteins with similar folds or partial sequence similarity can have different biological functions(24). Therefore, GNAT annotation alone is insufficient to predict STC resistance and functional validation is also required to identify enzymes with STC activity.

Functional metagenomics is well suited to this problem because it identifies DNA fragments through an expressed phenotype rather than through similarity to a known resistance sequence. Metagenomic or environmental DNA is cloned into an expression host (here, plasmids with constitutive promoters in *E. coli*) and grown in the presence of an antibiotic at a concentration above the host organism’s minimal inhibitory concentration (MIC). Clones that overcome inhibition and grow into colonies allow sequencing to directly link resistance to a metagenomic DNA fragment. This approach can identify novel resistance genes with low similarity to known references without culturing the original organisms(25, 26) (**Fig. 2**).

**Figure 2.**
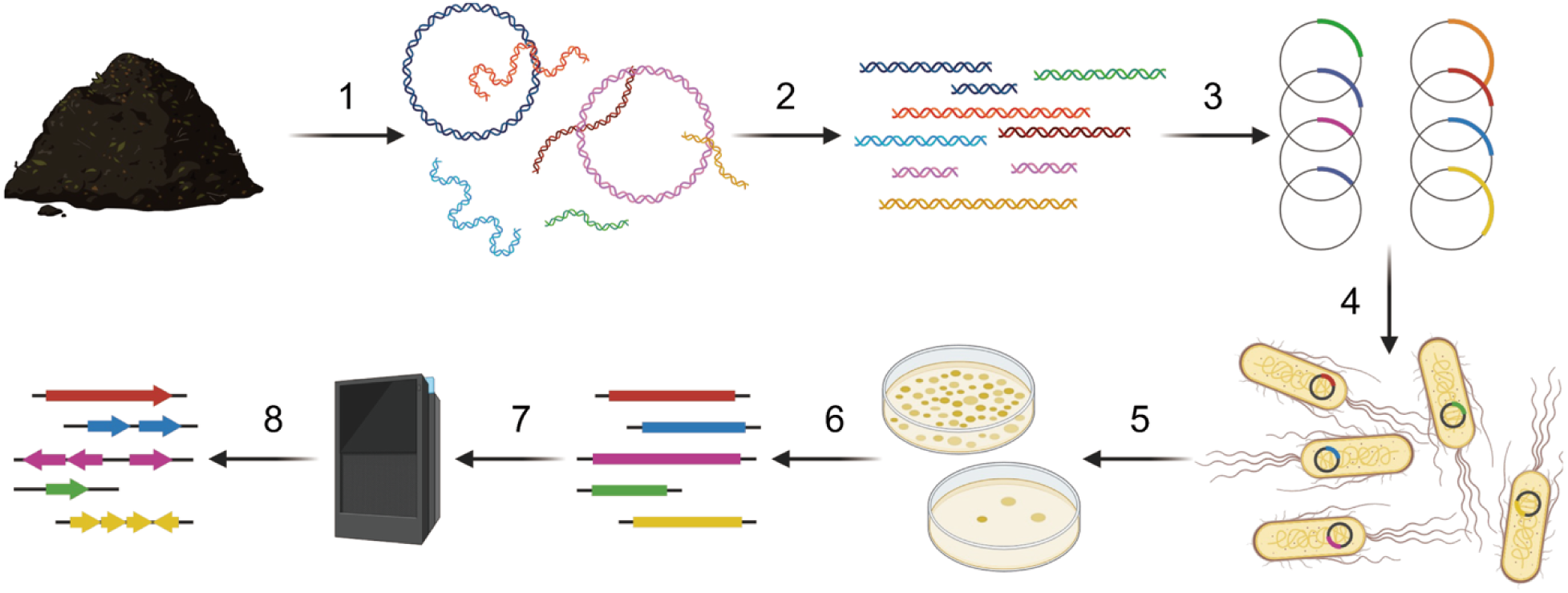
Functional metagenomic library construction. The general steps in the preparation and application of a functional metagenomic library are as follows: (1) Extraction of metagenomic DNA from a microbiome. (2) Fragmentation of metagenomic DNA to the preferred size range. (3) Cloning of inserts into expression vectors. (4) Transformation of host cells with vector library. (5) Screening or selection of a functional metagenomic library for a phenotype of interest. (6) Collection of screened or selected metagenomic fragments. (7) Sequencing of selected inserts. (8) Gene calling and annotation to identify potential genes underlying phenotypes of interest. Figure adapted from Allman *et al.*, 2025 (21), CC BY 4.0.

Our method for preparing functional metagenomic libraries, Mosaic Ends Tagmentation (METa) assembly, significantly improves library preparation by using Tn5-mediated tagmentation and homology-based assembly to increase cloning efficiency and reduce the amount of DNA required for library construction(21, 27). The (on average) 80-fold increase in library preparation efficiency allowed us to assemble a soil functional metagenomic library that captured ∼700 Gb of metagenomic DNA in previous work(27). This library uniquely captures sufficient metagenomic DNA to allow us to theoretically screen almost the entire functional diversity of even highly complex ecosystems such as those found in soil microbiomes, a process we refer to as deep functional metagenomics.

Here, we performed a deep functional metagenomic selection for NTC resistance on this large soil library. We analyzed 495,942 sequencing reads from an estimated 27,000 resistant colonies to identify candidate resistance determinants, focusing for now on acetyltransferase resistance mechanisms. By applying an abundance-weighted analysis pipeline (**Fig. 3**), we identified 265 acetyltransferase-like proteins encoded on 261 distinct PacBio circular consensus sequence inserts (CCS) from the top 5% of abundance-ranked representative ORF clusters. We selected 24 phylogenetically diverse candidates for experimental validation, obtained 18 sequence-verified constructs, and found that 10 conferred significant NTC-resistance when expressed in *E. coli*. Phylogenetic analysis placed these active proteins across multiple taxonomic branches, and we identified three new SAT lineages which we designate SatC, SatD, and SatE. We further tested these *sat*-expressing constructs against 4,6-disubstituted 2-deoxystreptamine (4,6-DOS) aminoglycosides and identified one construct that also conferred significantly decreased susceptibility to the antibiotic and WHO essential medicine amikacin. Together, these findings dramatically expand the known diversity of SAT resistance genes and provide new candidates for defining the broader streptothricin resistome.

**Figure 3.**
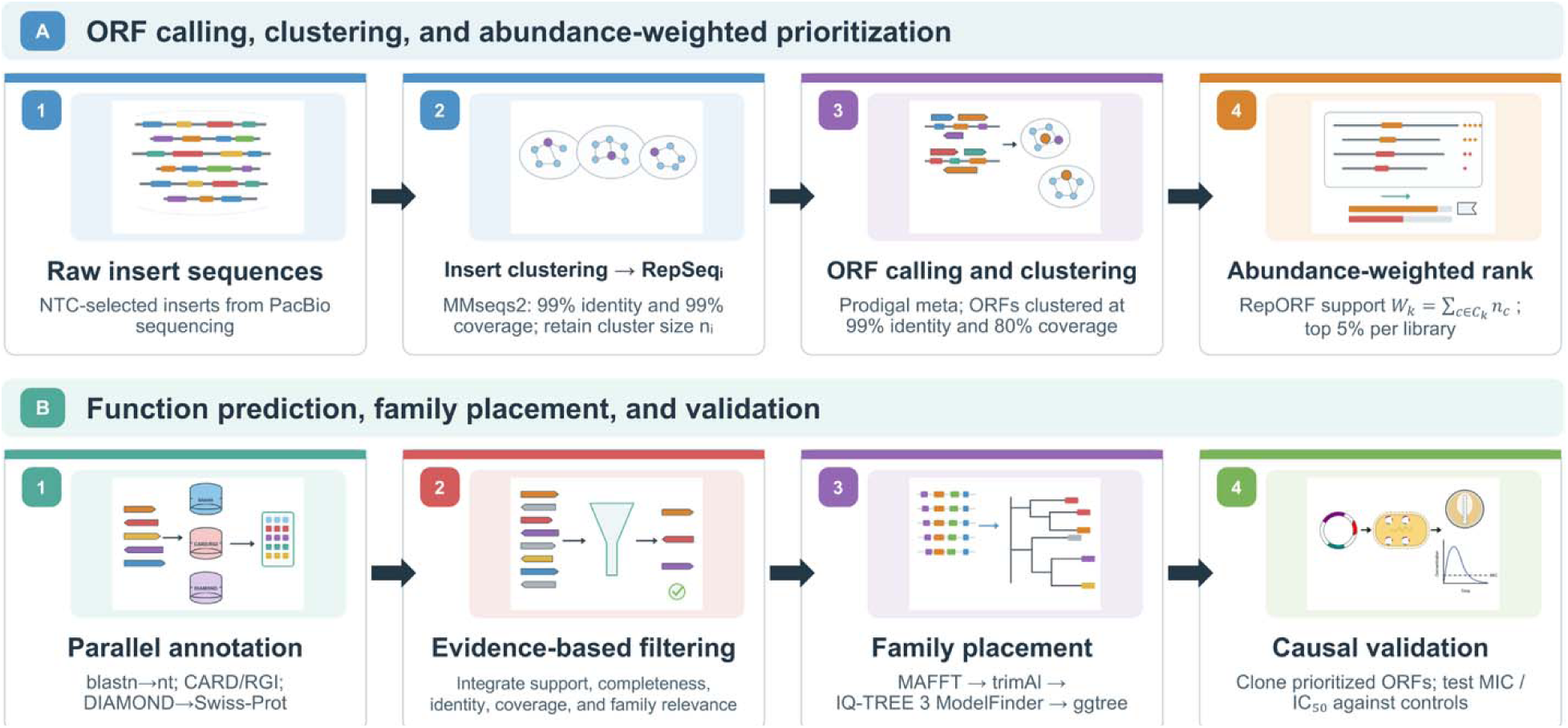
Workflow for abundance-weighted identification and annotation pipeline of nourseothricin-selected functional metagenomic libraries. (A) ORF calling, clustering, and abundance-weighted prioritization. Raw insert sequences were clustered with MMseqs2 at 99% nucleotide identity and 99% coverage to generate representative insert sequences (RepSeqs) while retaining the number of raw inserts represented by each cluster (*n_i_*). ORFs were predicted from RepSeqs using Prodigal in metagenomic mode and subsequently clustered at 99% nucleotide identity and 80% target-sequence coverage to generate representative ORF (RepORF) clusters. For each RepORF cluster j, abundance-weighted support was calculated as *W_k_* =∑*_c∈Ck_ n_c_*, where *W_k_* is the abundance-weighted support of RepORF cluster k,*C_k_* denotes the set of unique insert clusters contributing at least one member ORF to RepORF cluster k, c indexes an individual insert cluster within *C_k_*, and *n_c_* is the number of raw insert sequences represented by insert cluster c, and the top 5% within each library were prioritized for downstream analysis. (B) Function prediction, family placement, and experimental validation. Prioritized RepORFs were annotated in parallel using BLASTN against the nucleotide database, CARD/RGI for antimicrobial resistance determinants, and DIAMOND against Swiss-Prot. Candidates were filtered by integrated evidence including abundance support, sequence completeness, identity, coverage, and functional-family relevance. Selected candidates were aligned with reference sequences using MAFFT, trimmed with trimAl, and subjected to maximum-likelihood phylogenetic analysis using IQ-TREE 3; trees were visualized in ggtree. Prioritized ORFs were cloned and experimentally tested for resistance phenotypes.

## MATERIALS AND METHODS

### Nourseothricin functional metagenomic selection

The soil functional metagenomic library used in this study was constructed previously using METa assembly (27). Metagenomic DNA from soil gathered in Evanston, IL was tagmented using purified Tn5 transposase loaded with mosaic end oligos. Fragmented DNA was size-selected by gel extraction to target 1.5 kb to 5 kb, purified, and assembled into linearized pZE21-ME expression vector. We performed assembly across two reactions with one scaled up seven-fold compared to the other to give sub-libraries of different sizes. The resulting two pools of recombinant plasmids were introduced into *E.coli* DH10B (New England Biolabs, C3020K) by electroporation to make two soil functional metagenomic sub-libraries which we grew overnight in 50 ml of LB media (Miller) with 50 μg/ml kanamycin (VWR Life Science, 75856-68), concentrated to 10 ml in LB with 15% glycerol, and stored as 1 ml aliquots at -80°C. The detailed protocol has been described previously(27).

For NTC selection, we first determined the MIC (Minimum Inhibitory Concentration) of NTC against *E. coli* DH10B. NTC (nourseothricin sulfate, Dot Scientific, DSN51200-1) was prepared as a 100 mg/ml solution in water, sterilized by 0.22 μm filtration, and stored at -20°C until needed. Mueller-Hinton II cation adjusted (BD BBL, 212322) agar plates were prepared to contain 50 μg/ml kanamycin (MH+KAN) and NTC (0.5 μg/ml to 2048 μg/ml). The following *E. coli* strains were prepared at a density of 0.5 McFarland units and spread on to the agar plates and incubated overnight at 37°C: *E. coli* DH10B with an empty pZE21-ME plasmid, *E. coli* BW25113 strains with pGDP-expressed *rmtD* 16S rRNA methyltransferase, *ermC* 23S rRNA methyltransferase, and *stat* STC acetyltransferase (the three BW25113 strains came from the minimal antibiotic resistance platform [ARP], which was a gift from Gerard Wright) (Addgene kit no. 1000000143) (28).

Growth of the sensitive *E. coli* strains was inhibited by the presence of 2 μg/ml nourseothricin after overnight incubation; thus, we moved forward with a selection concentration of 4 μg/ml. To perform the deep functional metagenomic selection, frozen aliquots from the paired soil functional metagenomic sub-libraries were thawed on ice. A 1 ml aliquot of the ∼530 Gb soil library and 0.5 ml of ∼160 Gb soil library were combined and pelleted at 4,000 × g for 2 min and resuspended in 1.4 ml of Mueller-Hinton (MH) broth. The suspension was distributed at 100 µl per plate across 14 MH agar plates containing 50 µg/ml kanamycin and 4 µg/ml NTC. In parallel, an equivalent density lawn of *E. coli* DH10B carrying the empty pZE21-ME plasmid was plated on a single MH+KAN+NTC plate as an NTC-sensitive control. Plates were incubated at 37°C for 20 hr. Following incubation, NTC-resistant colonies were recovered from each plate by two washes with 750 µl of MH+KAN and the resulting suspensions were pooled. Plasmid DNA was isolated from two 1 ml aliquots of the pooled suspension using Monarch Spin Plasmid Miniprep Kit (NEB, T1110L). The remaining cells were pelleted at 4,000 × g for 7 min, resuspended in 1.3 ml of LB containing 15% glycerol, and stored at −80°C. Pooled plasmid DNA was subsequently used for Pacbio sequencing.

### Sequencing metagenomic DNA fragments that confer NTC resistance

Resistance-conferring metagenomic DNA fragments were sequenced by first amplifying them from the pooled plasmid miniprep described above. PCR (100 μl) contained 7 ng of miniprep DNA, forward and reverse primers at 500 nM (**Supplemental Table 1**), and Q5 polymerase Master Mix at 1X concentration (New England Biolabs, M0494S). Amplification occurred using a 2-step PCR protocol that alternated between 98°C for 10 sec and 72° for 4 min for 16 cycles. The amplicons were purified as before, quantified, and sent to the Roy J. Carver Biotechnology Center at the University of Illinois Urbana-Champaign for sequencing using a Pacific Biosciences (PacBio) SMRT Cell 8M Sequel II platform. Circular consensus sequence (CCS) analysis was performed using SMRTLink V8.0 with the following parameters: ccs –min-length 500 –max-length 12000 –min-passes 3 –min-rq 0.99. The resulting read were demultiplexed with lima (Pacific Biosciences) using default parameters.

### Insert clustering, representative ORF calling, functional annotation, and acetyltransferase candidate identification

PacBio CCS reads were clustered using MMseqs2(29) at 99% nucleotide identity and 99% alignment coverage of both the query and target sequences (--min-seq-id 0.99 -c 0.99 -- cov-mode 0 --cluster-mode 0). One representative insert sequence from each cluster (RepSeqs) was retained. Open reading frames (ORFs) were predicted on the RepSeqs inserts using Prodigal(30) in metagenomic mode. The ORF nucleotide sequences were subsequently clustered with MMseqs2 at 99% nucleotide identity and 80% target sequence coverage using greedy incremental clustering (--min-seq-id 0.99 -c 0.8 --cov-mode 1 --cluster-mode 2). The representative sequence from each cluster was designated as the representative ORF (RepORF). To estimate the raw-insert support associated with each RepORF cluster, member ORFs were mapped to their parent RepSeqs and subsequently to the corresponding insert-level MMseqs2 clusters. Each RepORF cluster was assigned an abundance-weighted support value based on the sizes of its contributing insert clusters calculated as

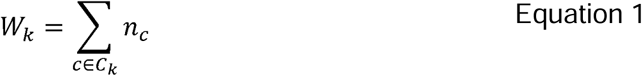

where *W_k_* is the abundance-weighted support of RepORF cluster *k*,*C_k_* denotes the set of unique insert clusters contributing at least one member ORF to RepORF cluster *k*, *c* indexes an individual insert cluster within *C_k_*, and *n_c_* is the number of raw insert sequences represented by insert cluster *c*. Each insert cluster contributed its size only once to a given RepORF cluster. RepORFs were ranked across the whole dataset according to their abundance-weighted cluster sizes, and the top 5% were retained for downstream analysis (**Fig. 3A**).

The selected RepORFs and their associated member ORFs were evaluated using two complementary protein annotation approaches. Antibiotic resistance gene (ARG) associated sequences were identified using Resistance Gene Identifier (RGI) v6.0.8 with the Comprehensive Antibiotic Resistance Database (CARD) v4.0.1(31). In parallel, predicted proteins were searched against UniProtKB/Swiss-Prot using DIAMOND BLASTp v2.2.2(32, 33). Annotation results are aggregated at the RepORF cluster level, while member-level evidence for individual ORFs is also retained. RepORF clusters supported by either annotation as encoding an acetyltransferase or acetyltransferase-like protein were retained as candidates for subsequent phylogenetic analysis (**Fig. 3B**).

### Phylogenetic analysis

For phylogenetic analysis of the putative acetyltransferase-like proteins, CARD amino acid sequences for the ontology term “acylation of antibiotic conferring resistance,” were downloaded. This set includes amino acid sequences for streptothricin, aminoglycoside, apramycin, virginiamycin, capreomycin, and chloramphenicol acetyltransferase enzymes. We also downloaded the amino acid sequence for the NAT nourseothricin acetyltransferase(34), SatB(21), and SrStat1, SrStat2(23). Multiple sequence alignment was performed with MAFFT v7.526 using automatic strategy selection (--auto)(35). Alignment trimming was evaluated with trimAl (36) at gap thresholds of 0.1, 0.2, 0.3, and 0.5.

Maximum-likelihood (ML) phylogenetic inference was performed with IQ-TREE v3.1.2(37). ModelFinder selected the best model from multiple protein substitution models according to the Bayesian information criterion. Tree searches were initiated from 20 parsimony trees (--ninit 20), with the five best initial trees subjected to maximum-likelihood optimization (-- ntop 5). Searches were terminated after 10 consecutive unsuccessful iterations without improvement in tree likelihood (--nstop 10). Branch support was estimated with 1,000 SH-like approximate likelihood ratio test (SH-aLRT) replicates and 1,000 ultrafast bootstrap replicates (--alrt 1000 -B 1000). The tree was visualized in R using ggtree(38) using custom scripts.

### Candidate selection, cloning, and construct sequencing

We selected 24 acetyltransferase-like RepORFs for further experimental evaluation.

Candidate ORFs were amplified by PCR using primer pairs carrying overlapping sequences for cloning assembly into the multiple cloning site of the pZA11 plasmid (39). Full primer sequences for target ORFs and pZA11 linearization are provided in **Supplemental Table 1**. PCR amplification was performed using Q5 polymerase Master Mix according to manufacturer’s recommendations (NEB, M0492S). Annealing temperatures were defined by the specific primer sequences, and extension times were adjusted to amplicon length in accordance with the manufacturer’s recommendations. Products of the expected size were separated on 1.5% 1X Lithium Acetate Borate (LAB) agarose (40). Bands were excised with a clean razor and purified using the Monarch Spin DNA Gel Extraction Kit (New England Biolabs, T1120L) according to the kit protocol.

Purified amplicons were combined at an approximately 2:1 insert-to-vector molar ratio in NEBuilder HiFi DNA Assembly Master Mix (New England Biolabs, E2621L) and incubated at 50°C for 15 min. Assembly products were purified using laboratory prepared SPRI beads(41, 42). The purified assemblies were introduced into chemically competent *E. coli* DH10B cells by heat-shock transformation, with incubation at 4°C for 30 min, followed by heat shock at 42°C for 30 s and cooling at 4°C for 2 min. Transformants were recovered in prewarmed SOC medium (New England Biolabs, B9020S) for 1 hr at 35°C and plated on LB agar containing 100 µg/ml ampicillin (Ward’s science, 470024-776). After overnight incubation at 35°C, individual colonies were inoculated into LB containing 100 µg/ml ampicillin and cultured at 35°C with shaking at 250 rpm overnight. Plasmid DNA was isolated using the Monarch Spin Plasmid Miniprep Kit.

Recovered plasmids were submitted to Plasmidsaurus for whole-plasmid long-read sequencing. Sequence records were compared with the expected pZA11 backbone and candidate ORF. ORFs that failed two amplification attempts (4 ORFs) and ORFs that showed serious errors upon sequencing (2 ORFs lacking stop codons) were excluded from further study.

### Taxonomic profiling of acetyltransferase-containing inserts

Taxonomic profiling was performed using protein-coding ORFs predicted from the 261 unique CCS inserts containing acetyltransferase-like candidates. A reference protein database was constructed from bacterial and archaeal species-representative genomes in Genome Taxonomy Database release R232 (GTDB R232)(43). For each named genus, a single representative genome was retained; for unclassified lineages, one representative genome was kept for each corresponding family, order, or phylum. Predicted proteins were searched against the reference database using MMseqs2 (sensitivity 7.5), with an E-value threshold of 1 × 10⁻, minimum sequence identity of 30%, and minimum query coverage of 50% (-e 1e-5 --min-seq-id 0.30 -c 0.50 --cov-mode 2). Taxonomic assignments were determined using a lowest common ancestor (LCA) approach.

Taxonomic origins of CCS inserts were inferred from non-target ORFs, excluding the acetyltransferase-like candidates. Consensus phylum assignments required either a single resolved non-target ORF or agreement among at least two non-target ORFs representing ≥60% of resolved assignments. Inserts not meeting these criteria were designated as unclassified.

Assignment confidence was categorized as high (≥3 supporting ORFs with ≥75% agreement), medium (≥2 supporting ORFs with ≥60% agreement), or low (a single resolved ORF). Class- and order-level assignments were determined hierarchically using the same criteria. Taxonomic assignments were mapped onto the acetyltransferase phylogeny and visualized using the R package ggtree(38), with annotation rings indicating phylum, class, order, and assignment confidence.

### Functional neighborhood Gene Ontology (GO) profiling

Protein-coding ORFs from the acetyltransferase-containing inserts were analyzed using Pfam and eggNOG-mapper for functional annotation. Protein domains were identified using HMMER v3.4 hmmscan(44) against Pfam release 38.2(45), with model specific Pfam gathering thresholds (--cut_ga). Pfam accessions were linked to InterPro entries using the official integration mappings from InterPro release 109.0(46). Gene ontology (GO) annotations were assigned using a custom script that combined GO terms from the InterPro2GO mapping distributed with InterPro release 109.0 and the Pfam2GO mapping obtained from the Gene Ontology website(47, 48). The proteins were also independently annotated with eggNOG-mapper v2.1.15 and the eggNOG v5.0.2 database(49). Homology searches were performed with DIAMOND in sensitive iterative mode, using an E-value threshold of 1×10^-3^, and a minimum query coverage of 50%. GO transfer used the non-electronic evidence setting (--go_evidence non-electronic). GO annotations independently supported by Pfam/InterPro mappings were retained regardless of the eggNOG filter. Retained annotations were merged and deduplicated for each ORF. GO identifiers were normalized against the GO Basic ontology release dated 26 July 2026 and mapped to generic GO-slim terms. Acetyltransferase-like target ORFs were excluded from the functional neighborhood analysis. GO-slim categories were summarized as binary presence or absence for each CCS insert, with each category counted once per insert.

The 24 inserts corresponding to the candidates selected for cloning were additionally annotated using Bakta v1.12.0 with the full database v6.0(50). The run used Pyrodigal v3.7.1 for coding-sequence prediction(51), AMRFinderPlus v4.2.7 for antimicrobial resistance annotation(52), PyHMMER v0.12.3 for profile-HMM searches(53), DIAMOND v2.2.6, and BLAST+ v2.17.0. Inserts were analyzed as short environmental contigs, with coding sequences predicted in metagenome mode. Gene annotations were primarily based on the RGI and UniProtKB/Swiss-Prot results described above and were supplemented with Bakta annotations. Gene schematics for all 24 CCS inserts were manually generated in Microsoft PowerPoint based on the combined annotation results.

### NTC antimicrobial susceptibility testing by agar dilution

The 18 successfully cloned candidate constructs (out of 24 attempted) were evaluated for NTC susceptibility by agar dilution, with the empty pZA11 vector strain included as negative control (54). MH agar containing 100 µg/ml ampicillin (MH+AMP) was supplemented with NTC at final concentrations of 0 μg/ml and 0.5 to 1,024 µg/ml by 2-fold jumps. Colonies of clones to be tested were suspended in MH broth and adjusted to a 0.5 McFarland standard, corresponding to approximately 1×10^8^ CFU/ml. The adjusted suspensions were diluted 10-fold in MH broth, following which 1 µl of each diluted suspension, corresponding to an inoculum of approximately 1×10^4^ CFU, was transferred on to each agar plate beginning with the 0 μg/ml plate and moving up the dilution gradient. A 48-pin replicator (Boekel Scientific, 140501) was used for parallel inoculation and was sterilized between plates with 70% ethanol followed by flaming. Inoculated plates were incubated at room temperature until the spots were absorbed, then plates were inverted and incubated at 35°C for 16–20 h. The agar dilution MIC was defined as the lowest NTC concentration preventing visible growth.

### Nourseothricin and aminoglycoside susceptibility testing by broth microdilution assay

Broth microdilution assays were performed for hits 8, 57, 178, 185, 259, and the empty-vector control pZA11 against nourseothricin, amikacin (Thermo Scientific, J63862.14), kanamycin, and spectinomycin (Thermo Scientific, J66034.06)(54). Two-fold serial dilutions of each antibiotic were prepared in MH+AMP broth. Final antibiotic concentrations were 0.5 to 256 µg/ ml for NTC, 0.032 to 16 µg/ ml for amikacin, 0.032 to 16 µg/ml for kanamycin, and 1 to 512 µg/ml for spectinomycin. Fresh colonies were suspended and adjusted to 0.5 McFarland, followed by a 100-fold dilution in MH+AMP broth. An equal volume (50 µl) of the diluted bacterial suspension was added to 50 µl of the antibiotic dilution in 96-well plates, giving a final volume of 100 µl and a target inoculum of approximately 5×10 CFU/ml. Growth controls without antibiotic and uninoculated sterility controls were also included. The broth microdilution assays were conducted in two experiments. In the first experiment, hits 57, 178, 185, 259, and the empty-vector control pZA11 were tested against nourseothricin and amikacin. In the second experiment, hits 8, 57, 178, 185, 259, and pZA11 were tested against kanamycin and spectinomycin, and hit 8 and pZA11 were additionally tested against nourseothricin and amikacin. Each strain–antibiotic combination was tested with four replicates. Plates were sealed with a Breathe-Easy membrane (MilliporeSigma, Z380059) and incubated at 35°C with shaking for 16–20 h. After subtracting blank absorbance values, bacterial growth was recorded as OD_600_ with a 1 cm pathlength correction using an Epoch 2 plate reader (BioTek), and the concentration-response curves were fitted to four-parameter Hill equations in R package drc (v3.0)(55) to determine 50% inhibitory concentration (IC_50_) values. For nourseothricin and amikacin, concentration-response curves from the two experiments were plotted separately, while IC_50_ values from both experiments were retained in the combined IC_50_ analysis. Differences between IC_50_ values for control E. coli and the tested clones were evaluated for significance with multiple comparison correction using the Dunnett test in R package multcomp (v1.4)(56).

### Protein structure prediction and visualization

Protein structures were predicted for candidate proteins using AlphaFold 3 through the AlphaFold Server(57). Each protein was submitted separately as a single protein chain. Five models were returned per submission. The model with the highest-ranking score was selected for subsequent analysis. Local prediction confidence was represented using the predicted local distance difference test (pLDDT). GNAT-family regions were annotated using HMMER hmmscan against Pfam, applying model-specific gathering thresholds (--cut_ga).

Structural alignments and visualization were performed in UCSF ChimeraX v1.12(58). Each candidate model and a SatB model(21) were independently superimposed against the SatA crystal structure (3PP9)(20, 59) and an aminoglycoside acetyltransferase AAC(6’)-Ib crystal structure (2QIR) (60) using MatchMaker. Root-mean-square deviations (RMSDs) were reported both for the retained residue pairs and for all initially matched pairs under the final superposition, together with the corresponding pair counts. Following superposition, SatA was hidden when visualizing each target, and a common viewing orientation was maintained. Cartoon representations were colored by pLDDT, while purple and green outlines marked the first and second annotated GNAT regions, respectively. Panels were independently centered and scaled for display.

### Statistical analysis and data visualization

Statistical analyses and data visualization were performed in R version 4.4.3 with ggplot2 3.5.1. All analysis pipeline and custom code are available at github. (https://github.com/croftsmicrolab/arg-annotator, https://github.com/croftsmicrolab/Soil_NTC_paper)

## RESULTS

### Nourseothricin resistance selection and sequencing

To better characterize the soil streptothricin resistome, we performed a functional metagenomic selection for NTC resistance using a combined soil library encompassing ∼700 Gb of metagenomic DNA. Plating of this library resulted in the capture of an estimated ∼27,000 NTC resistant colonies with a range of colony sizes (**Supplementary Fig. 1A**). After collection of the surviving colonies by combining them together into a single aliquot, we extracted resistance-conferring plasmids, amplified, and sequenced the captured metagenomic DNA fragments. After sequence preprocessing, the dataset comprised 495,942 raw insert sequences. The average read length was 1.846 ± 0.525 kb, in general agreement of our previously reported average insert size of 1.94 kb for the whole library + 500 bp of plasmid backbone (**Supplementary Fig. 1B**) (27).

### Abundance-weighted ORF clustering prioritizes 265 acetyltransferase-like candidates for phylogenetic analysis

Clustering of the raw insert sequences by nucleotide identity generated 126,158 insert clusters, including 107,192 singletons. One representative insert sequence was retained from each cluster for subsequent ORF prediction (**Supplementary Fig. 2**). We used an abundance weighted analysis approach to select targets for further study from this population (**Fig. 3**). A total of 354,401 ORFs were predicted from the RepSeqs (giving an average of 2.81 ORFs per clustered insert) and further clustered into 281,926 RepORF clusters. After applying abundance-weighted support value calculation, there were 237,496 singletons (**Supplementary Fig. 3**) (**Fig. 3A**). The top 5% of the ranked distribution was retained for downstream analysis, yielding 14,097 RepORFs. ARG-related evidence was detected in 22,516 member ORFs distributed among 5,701 RepORF clusters. Annotation-based screening of the 14,097 selected RepORFs identified 265 acetyltransferase-like candidate proteins originating from 261 distinct CCS inserts (**Fig. 3B**).

To examine the phylogenetic relationships of the candidate acetyltransferases, we constructed an unrooted phylogenetic tree using the 265 candidate RepORFs together with 88 reference antibiotic resistance acetyltransferases from CARD (353 proteins total; **Fig. 4**). Many candidate RepORFs were distributed within the broader phylogenetic region containing known streptothricin acetyltransferases but occupied relatively distant branches from the characterized SAT references.

**Figure 4.**
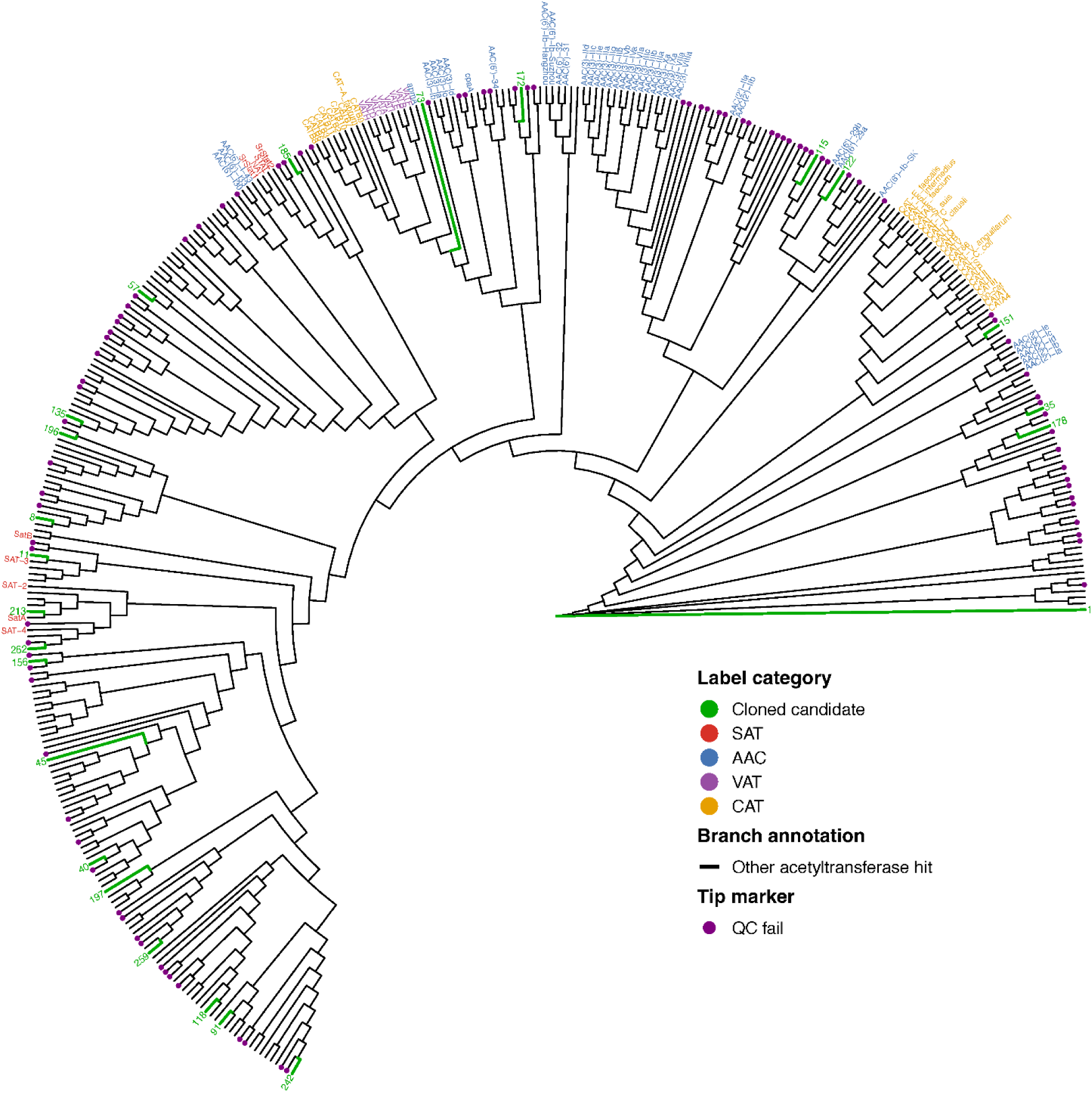
Phylogenetic distribution of candidate acetyltransferases relative to known acetyltransferases. An unrooted phylogenetic tree was constructed from 265 candidate RepORFs (black branches) and 88 reference acetyltransferases (353 proteins total). Candidate RepORFs are shown in green and reference proteins are colored by antibiotic resistance function. Candidates that failed subsequent sequence verification are shown in purple tip.

### Candidate acetyltransferases occur across diverse taxonomic and functional contexts

To further characterize the genomic context of these candidates, we examined the inferred taxonomy of the 261 candidate-containing inserts and predicted functions for genes syntenic to the targeted acetyltransferases. Putative phylum-level assignments were obtained for 209 of 261 CCSs (80.1%), with *Chloroflexota* representing the largest assigned group, followed by *Bacillota*, *Actinomycetota*, and *Acidobacteriota* (**Fig. 5B, Supplementary Fig.4**). Functional annotation of neighboring ORFs identified a broad range of GO-slim categories associated with the candidate-containing inserts (**Supplementary Fig. 5**). Among the molecular function categories, transferase and hydrolase activities were the most frequently represented, while biological process annotations included diverse metabolic, regulatory, and DNA-associated functions.

**Figure 5.**
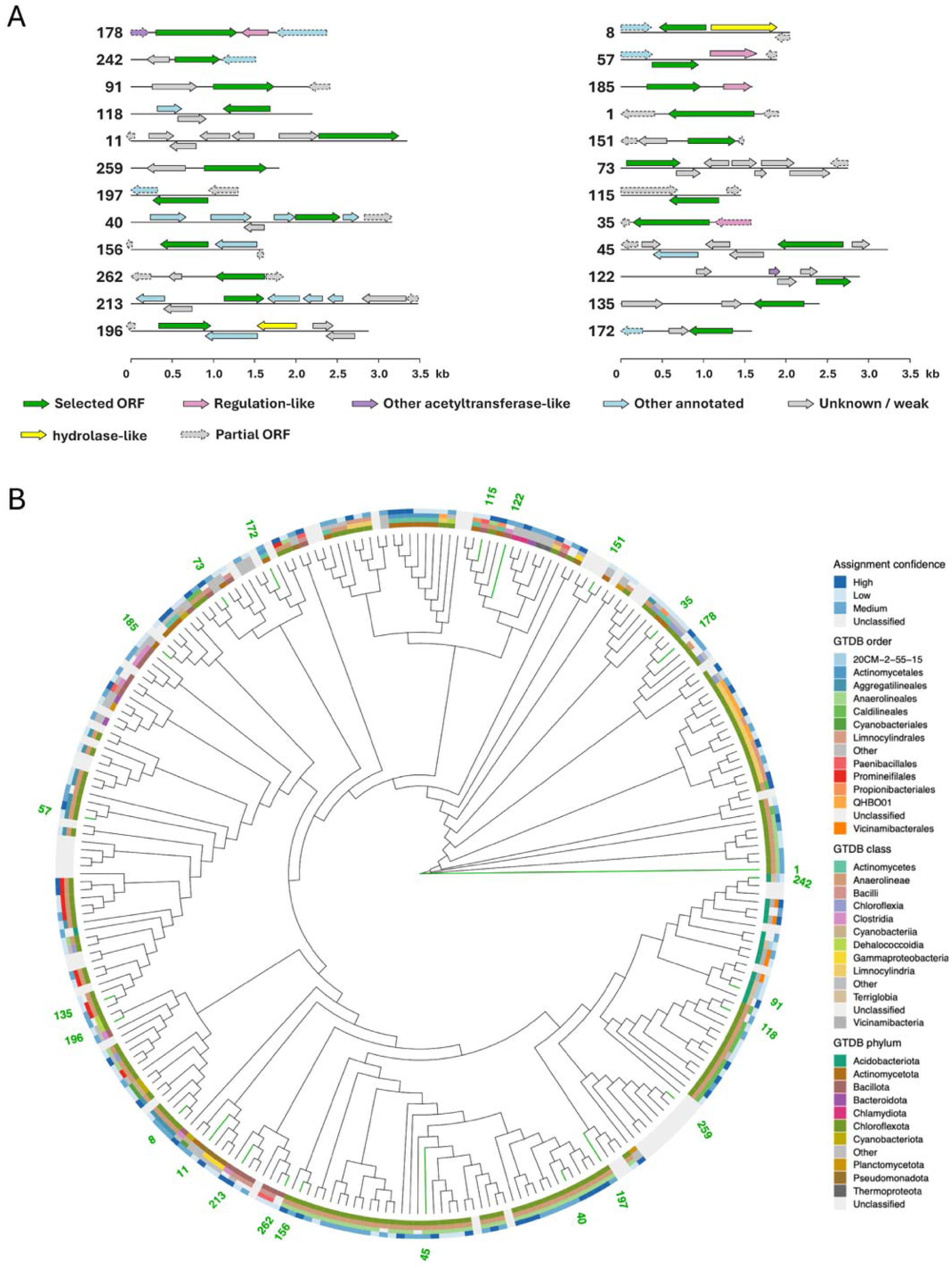
Genomic context and phylogenetic distribution of candidate acetyltransferases. (A) Gene schema of 24 candidates metagenomic DNA fragments. Arrows represent predicted ORFs, with arrow direction indicating coding orientation and arrow length corresponding to ORF length. Selected acetyltransferase-like ORFs are shown in blue. Neighboring ORFs are colored according to functional annotation: regulation-like (pink); other acetyltransferase-like (purple); hydrolase-like (yellow); other annotated proteins (light blue); unknown or weakly annotated proteins (gray). ORFs predicted to be incomplete are indicated by dashed outlines. The scale bar indicates insert length in kilobases (kb). (B) Distribution of taxonomic assignments across the candidate acetyltransferase phylogeny. Rings indicate GTDB phylum-, class-, and order-level assignments, together with assignment confidence. Green labels indicate candidate ORFs selected for further study.

### Identification and functional validation of diverse NTC acetyltransferases within new Sat phylogenetic lineages

Twenty-four candidate streptothricin acetyltransferase genes were selected for experimental validation based on the location of their corresponding predicted enzymes across our acetyltransferase phylogenetic tree (**Fig. 4**). The corresponding inserts showed diverse gene organizations surrounding the selected acetyltransferase-like ORFs (**Fig. 5A**), including other acetyltransferase-like, regulation-like, and hydrolase-like genes. Four candidate genes could not be amplified from the original soil slurry DNA, and two additional candidate genes were excluded after cloning resulted in inserts lacking stop codons, leaving 18 sequence-verified ORFs for functional testing (hit numbers 8, 35, 40, 57, 91, 118, 122, 135, 151, 156, 178, 185, 196, 197, 213, 242, 259, and 262). Comparison of the sequenced constructs with their original predicted ORFs showed some minor differences for several candidates. Therefore, all subsequent analyses used protein sequences translated from the experimentally verified constructs.

To determine if the selected ORFs were actual streptothricin acetyltransferases, we performed agar dilution assays to measure changes in STC susceptibility in *E. coli* expressing the genes. Of the18 constructs, 11 conferred detectable NTC resistance relative to the pZA11 empty-vector control (hits 178, 242, 91, 259, 197, 118, 40, 8, 156, 57, and 262) and 10 showed at least a 16-fold increase in MIC (all except hit 178) (**Fig. 6A**). Hits 8, 256, and 262 showed particularly high levels of resistance, with growth observed at NTC concentrations above 512 µg/ml. For comparison, the empty-vector control showed no detectable growth at 0.5 µg/ml NTC and hit 178 showed no growth past 1 μg/ml NTC.

**Figure 6.**
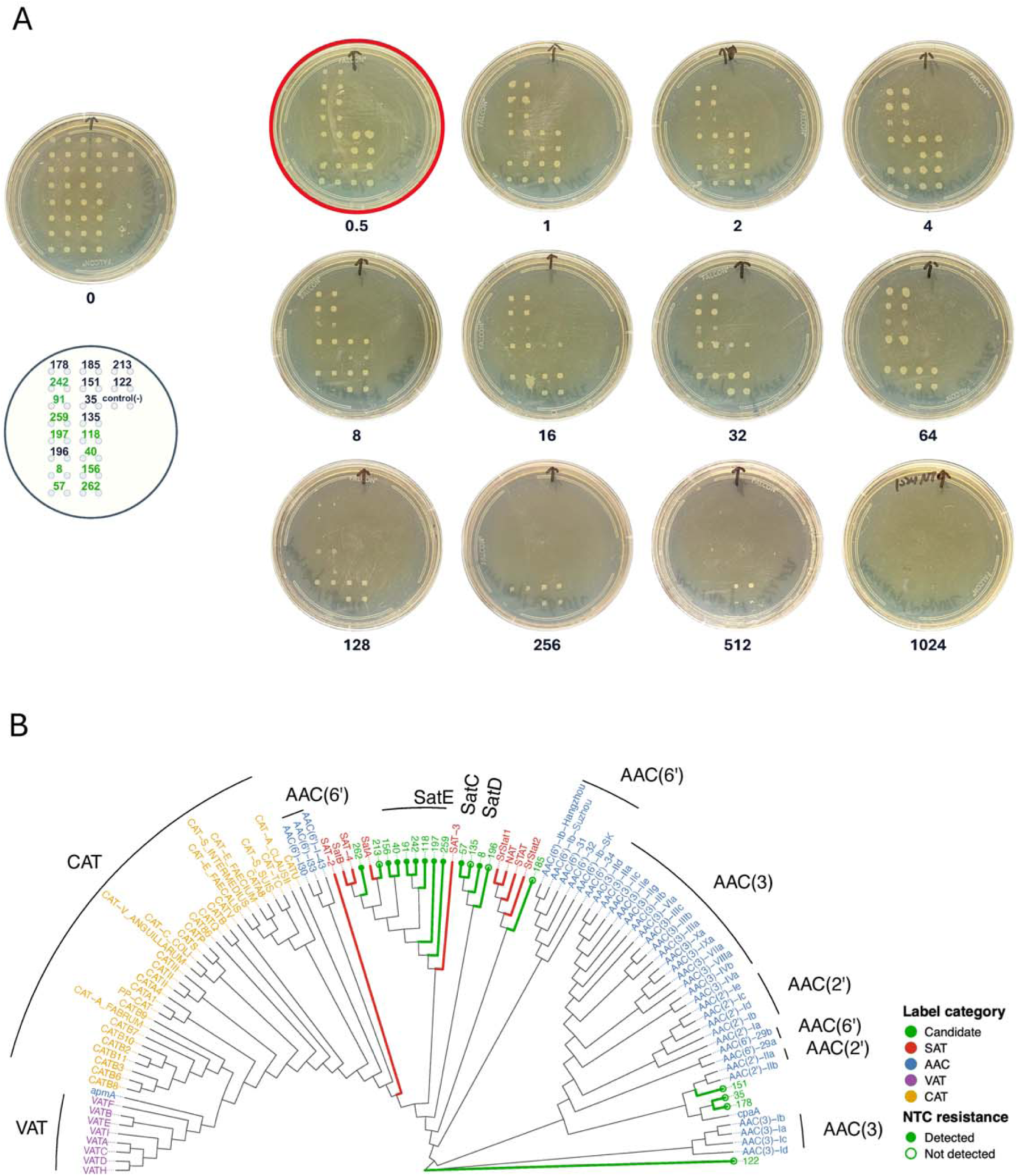
Cloning and functional validation identify phylogenetically diverse NTC resistance-conferring acetyltransferases. (A) Agar dilution assay of the 18 sequence-verified candidate constructs across increasing concentrations of nourseothricin (NTC; 0–1,024 µg/mL, 2-fold serial dilutions). Each agar plate represents a single NTC concentration, as indicated below the plate. Candidate constructs were spotted at fixed positions across all plates, with the plate layout shown in the schematic at left. Each construct was tested in duplicate. The pZA11 empty-vector strain was included as a negative control. Green labels indicate highly NTC-resistant constructs. The red circle indicates the lowest NTC concentration at which the empty-vector negative control exhibited no growth. (B) Sequence-resolved phylogenetic analysis of the 18 sequence-verified candidate proteins and 88 reference acetyltransferases. The unrooted tree was reconstructed using protein sequences obtained from the sequenced candidate constructs. Sequence-verified candidates are shown in green, SAT-related reference acetyltransferases in red, and other reference acetyltransferases in various colors. NTC resistance phenotypes determined by agar dilution are indicated by circles at the tips, with filled green circles indicating high levels of resistance and open green circles indicating no or negligible detected resistance. The three designated resistance-associated lineages, SatC, SatD, and SatE, are indicated on the tree.

We further determined the phylogenetic relationships of the 18 experimentally tested ORFs. Predicted amino acid sequences for the 18 hits were analyzed alongside the same 88 reference resistance acetyltransferases used in our earlier analysis, generating a 106 protein phylogenetic tree (**Fig. 6B**). NTC resistance phenotypes were mapped onto the reconstructed tree, revealing that resistance conferring candidates were distributed across multiple branches, including several branches distinct from previously characterized streptothricin acetyltransferases.

Pairwise amino acid sequence identities among the 18 candidates and between the 18 candidates and 88 reference proteins were further examined to assess their sequence relationships (**Supplementary Fig. 6)**. Based on these sequence relationships, together with their phylogenetic positions and experimentally confirmed NTC resistance (**Fig. 6A**), three additional resistance-associated lineages were provisionally designated as SatC (hit 57), SatD (hit 8), and SatE (hit numbers 40, 91,118, 156, 197, 242, 259). The seven SatE proteins form a relatively close branch, sharing 37.9–73.4% pairwise identity with one another (mean, 48.9%) (**Supplementary Fig. 6A**), while showing lower identity to known SATs (the highest observed identity was 47.5% between hit 156 and SatA) (**Supplementary Fig. 6B**). SatC (hit 57) and SatD (hit 8) were more divergent from characterized SATs and other candidates, with maximum identities of 31.0% to SrStat1 and 31.1% to SAT-3, respectively, and 33% to each other (**Supplementary Fig. 6**).

### *satC* expression confers significant NTC resistance and decreases susceptibility to amikacin and kanamycin

To explore whether expression of *satC, satD,* or *satE* conferred resistance to other antibiotics, five constructs were selected for quantitative broth microdilution analysis against amikacin, kanamycin, spectinomycin and NTC. Hits 57, 8, and 259 were selected as representatives of the SatC, SatD and SatE lineages, respectively. In addition, hits 178 and 185 were selected because of their phylogenetic proximity to aminoglycoside acetyltransferase (AAC) enzymes (**Fig. 6B**).

Broth microdilution against NTC reconfirmed the resistance phenotypes observed in the agar dilution assay (**Fig. 7A and Supplementary Fig. 7A**). The pZA11 empty-vector control had a mean IC_50_ of 0.88 ± 0.76 µg/ml. In comparison, the NTC dose-response curves for *satC* (hit 57) and *satE* (hit 259) were strongly shifted to the right (**Supplementary Fig. 7A**), with significantly higher IC_50_ values of 16.4 ± 1.24 and 71.7 ± 4.12 µg/ml, respectively (P_adj_ < 0.0001). *satD* (hit 8) showed the strongest NTC resistance phenotype, with 50% growth inhibition not reached at the highest NTC concentration tested (256 µg/ml) (**Supplementary Fig. 7A**), consistent with its strong resistance phenotype observed by agar dilution (**Fig. 6A**). Hit 178 showed a small increase in IC_50_ (2.51 ± 0.38 µg/ml), but this difference was not statistically significant compared to control (P_adj_ = 0.156). Hit 185 was not significantly different from the pZA11 control (P_adj_ = 0.999) and had a similar IC_50_ value (1.02 ± 0.07 µg/ml).

**Figure 7.**
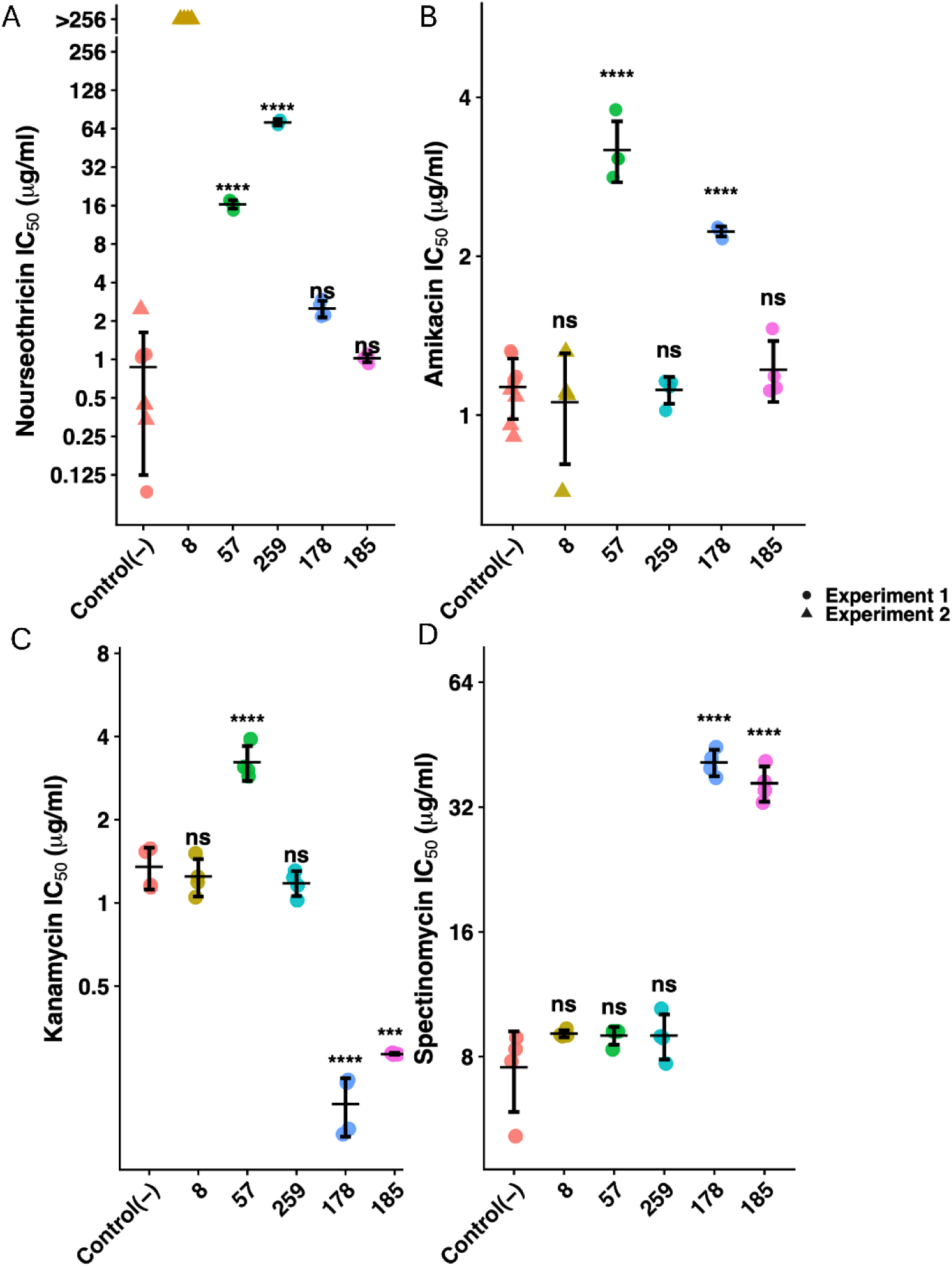
Antibiotic susceptibility profiles of candidate resistance clones. IC_50_ values for (A) nourseothricin, (B) amikacin, (C) kanamycin, and (D) spectinomycin. Points represent individual measurements, with circles and triangles indicating experiments 1 and 2, respectively. Error bars indicate mean ± SD. Statistical significance was determined using Dunnett’s test with the empty-vector control as the reference. ns, not significant; ****P < 0.0001.

Testing against amikacin and kanamycin revealed a distinct susceptibility profile for the *E. coli* clone expressing *satC* (hit 57) (**Fig. 7B and 7C, and Supplementary Fig. 7B and 7C**). Expression of *satC* (hit 57) increased the amikacin IC_50_ from 1.13 ± 0.15 µg/ml for the pZA11 control to 3.18 ± 0.42 µg/ml (∼2.8-fold; P_adj_ < 0.0001) and the kanamycin IC_50_ from 1.35 ± 0.23 to 3.23 ± 0.47 µg/ml (∼2.4-fold; P_adj_ < 0.0001). In contrast, *satD* (hit 8) and *satE* (hit 259) expression did not significantly alter susceptibility to either antibiotic. Expression of hit 178 also showed a significant increase in amikacin IC_50_ (2.23 ± 0.05 µg/ml; P_adj_ < 0.0001), but its kanamycin IC_50_ decreased significantly to 0.19 ± 0.04 µg/ml (P_adj_ < 0.0001). Expression of hit 185 similarly showed a decreased kanamycin IC_50_ (0.28 ± 0.002 µg/ml; P_adj_ < 0.0001), while its amikacin IC_50_ remained comparable to pZA11 (**Fig. 7B and 7C, and Supplementary Fig. 7B and 7C**).

Spectinomycin susceptibility was not altered by expression of any of the *sat* genes (hits 57, 8, and 259) (**Fig. 7D; Supplementary Fig. 7D**). Their mean IC_50_ values (8.99 ± 0.44, 9.09 ± 0.18, and 8.99 ± 1.12 µg/ml, respectively), were not significantly different compared to 7.54 ± 1.66 µg/ml for the pZA11 control. In contrast, expression of hits 178 and 185 showed pronounced increases in spectinomycin IC_50_ to 41.0 ± 2.99 and 36.5 ± 3.54 µg/ml, respectively (Padj < 0.001).

### Analyses of sequences and predicted structures reveal conserved functional features among divergent Sat acetyltransferases

To further examine the sequence and structural features of the identified NTC resistance-conferring acetyltransferases, we compared the predicted candidate proteins against previously characterized streptothricin acetyltransferases. Multiple sequence alignment showed that several conserved functional residues previously implicated in SatA-mediated streptothricin detoxification were also retained in our candidate proteins(20, 59) (**Supplementary Fig. 8**). These include the predicted catalytic residue Glu137 and residues associated with the putative streptothricin-binding pocket, including Leu136, Ala145, Tyr149, Phe154, and Tyr164 (SatA numbering). A notable exception was in SatD (hit 8), in which the residue corresponding to Asp160 of SatA is an asparagine (Asn) (**Supplementary Fig. 8**).

Comparison of the predicted structures of the candidate proteins with *B. anthracis* SatA and the aminoglycoside acetyltransferase AAC(6′)-Ib revealed both conserved and divergent structural features (**Fig. 8A**). Structural similarity was assessed using local structural RMSD, calculated from well-aligned residue pairs after iterative refinement, and overall structural RMSD, calculated across all matched residue pairs (**Fig. 8B**).

**Figure 8.**
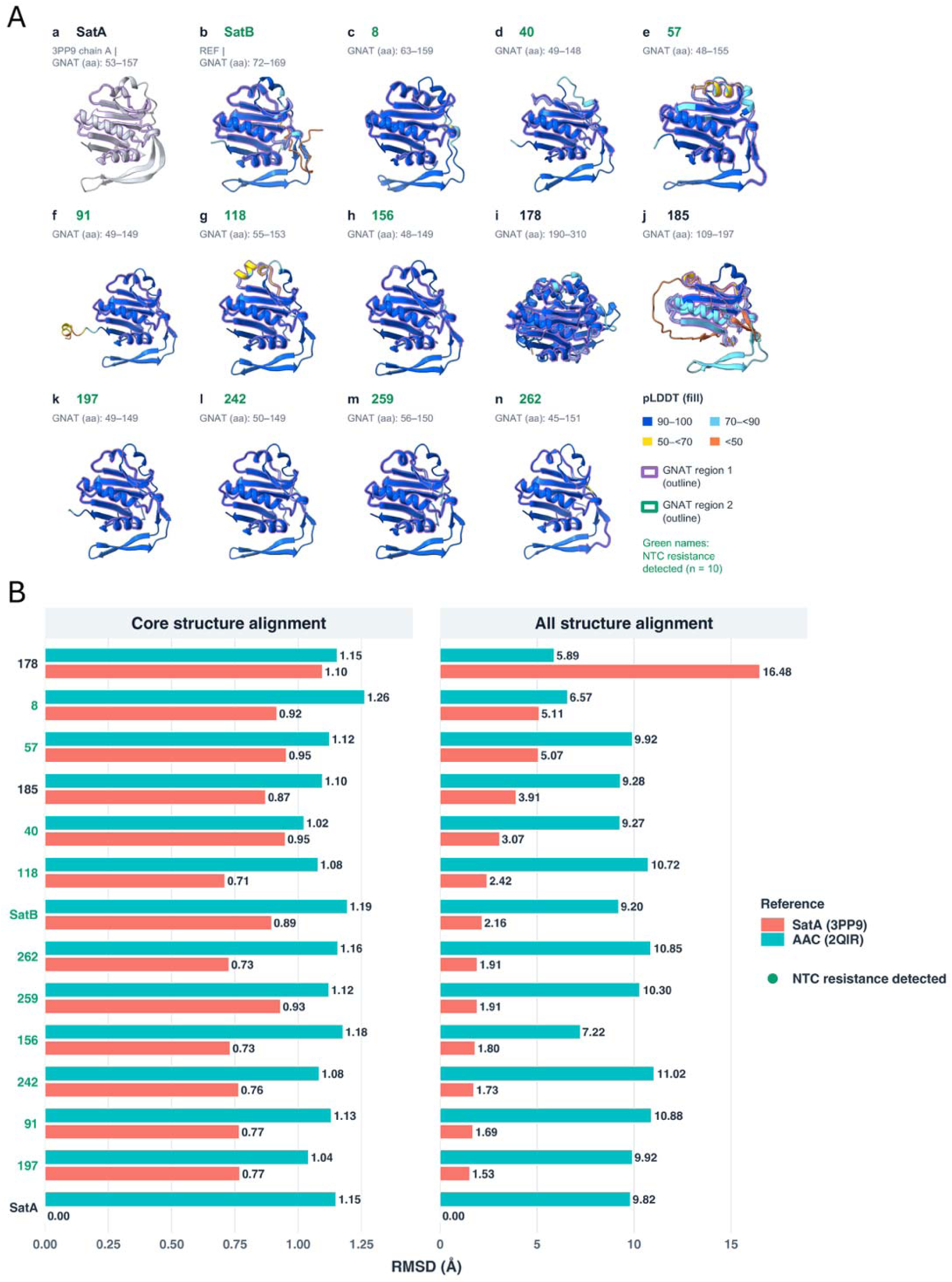
Sequence and structural comparison of candidate acetyltransferases with characterized acetyltransferases. (A) Structural superposition of the AlphaFold3-predicted candidate proteins and SatB onto the SatA structure. Predicted structures are colored according to pLDDT scores: dark blue, 90–100; cyan, 70–90; yellow, 50–70; and orange<50. The experimental SatA structure (PDB 3PP9) is shown in gray. GNAT regions are outlined in purple, with the second GNAT region outlined in green. Candidate names shown in red indicate constructs that conferred NTC resistance. (B) Structural comparison of candidate proteins with SatA and the distantly related aminoglycoside acetyltransferase AAC(6′)-Ib. RMSDs are shown for residue pairs retained after pruning and for all initially matched residue pairs under the final superposition. Pink bars indicate comparisons with SatA (PDB 3PP9), and teal bars indicate comparisons with AAC(6′)-Ib (PDB 2QIR).

Comparison with SatA revealed conserved structural regions across the candidate proteins, as indicated by low local predicted structural RMSDs (**Fig. 8B**) (*i.e.,* between 0.71 Å and 1.1 Å). Members of the SatE lineage showed particularly high structural similarity to SatA, with local RMSDs of 0.71 Å to 0.95 Å and overall RMSDs of 1.53 Å to 3.07 Å. In contrast, SatC (hit 57) and SatD (hit 8) showed greater structural divergence outside the well-aligned regions. Although their local predicted RMSDs remained low (0.95 Å and 0.92 Å, respectively), their corresponding overall RMSDs were substantially higher (5.07 and 5.11 Å, respectively; **Fig. 8B**).

Comparison with AAC(6′)-Ib further distinguished the structural similarities of these candidates. Most showed higher overall RMSDs to AAC(6′)-Ib than to SatA, including SatC (9.92 Å versus 5.07 Å) and SatD (6.57 Å versus 5.11 Å), indicating greater overall structural similarity to SatA. Additional structural variation was observed outside the conserved regions. In particular, hit 91, a member of the SatE lineage, contained an extended C-terminal region relative to SatA, whereas the previously characterized SatB contains an N-terminal extension (**Fig. 8A**). Together, these comparisons indicate that the newly identified Sat lineages retain conserved SatA-like structural regions despite substantial variation in their overall structural organization.

## DISCUSSION

Streptothricins are naturally produced antibiotics that are believed to be widespread in soil microbial communities, but the known streptothricin resistome remains remarkably limited compared to similar soil-derived antibiotics like the aminoglycosides (7). Only a few resistance mechanisms against streptothricins have been experimentally characterized, and the best characterized class, streptothricin acetyltransferases, only contains a small number of enzyme families, creating a substantial gap between the predicted ecological prevalence of streptothricins and their resistance (7). Here, the analysis of a large scale soil functional metagenomic dataset obtained through NTC selection reveals that the diversity of putative streptothricin resistance determinants is far greater than current representation in the literature suggests. We identified 265 acetyltransferase-like candidates distributed across a broad phylogenetic range and diverse taxonomic and functional genomic contexts. Experimental validation identified multiple previously uncharacterized proteins capable of conferring NTC resistance, including three phylogenetically distinct resistance-associated lineages provisionally designated SatC, SatD, and SatE.

An important contribution of this study is the use of abundance-weighted ORF clustering to prioritize resistance-associated sequences that would be difficult to identify through raw insert abundance alone. The NTC-selected dataset was sparse, with 107,192 of 126,158 raw insert clusters represented by singletons (84.97%) (**Supplementary Fig. 2**). Several experimentally validated NTC resistance determinants originated from singleton or low abundance insert clusters but were prioritized through aggregation of closely related ORFs across different insert clusters (**Supplementary Fig. 2 and 3**). By incorporating the abundance of contributing insert clusters into ORF-level ranking, this approach concentrated resistance-associated sequence signals that were dispersed across individual metagenomic fragments. Importantly, ten of the 18 sequence-verified candidates (55.6%) conferred at least a 16-fold increase in NTC MIC when individually expressed in *E. coli* (**Fig. 6A**). Together, these findings demonstrate the utility of abundance-weighted candidate prioritization for identifying functional resistance determinants within highly heterogeneous metagenomic datasets. Notably, this hit rate suggests that dozens to hundreds of additional NTC resistance genes remain among the untested candidates.

The experimental validation results also illustrate why resistance-associated inserts do not necessarily yield resistance when individual predicted ORFs are expressed separately. Because functional metagenomic selection identifies resistance-conferring DNA fragments rather than individual genes, the resistance phenotype of an original insert may arise from another ORF encoded on the same fragment, or a combination of ORFs. For example, hit 213 shared 89.3% amino acid identity with SatA but lacked a substantial N-terminal region and did not confer detectable NTC resistance. Although no other clearly identifiable streptothricin resistance determinants were found on its corresponding insert, the lack of resistance may reflect an incomplete or incorrectly annotated ORF or contributions from neighboring genes. Furthermore, hit 178 and hit 185 showed distinct changes in susceptibility to other antibiotics despite conferring little or no NTC resistance (**Fig. 7**), suggesting that some acetyltransferase-like candidates may have functions beyond streptothricin resistance. Automated annotation was also insufficient to capture these activities, as hit 178 and hit 185 lacked aminoglycoside resistance annotations, while experimentally validated NTC resistance determinants such as hit 57 and hit 118 lacked relevant Bakta annotations. Together, these findings highlight the limitations of relying on sequence similarity or automated annotation alone and emphasize the value of combining abundance-weighted prioritization, phylogenetic analysis, and individual gene validation to identify functional resistance determinants.

The inferred taxonomic distribution of the candidate-containing inserts further supports a broad environmental reservoir of streptothricin resistance. Candidate-containing inserts were assigned to multiple bacterial phyla, with *Chloroflexota* representing the largest group, followed by *Bacillota*, *Actinomycetota*, and *Acidobacteriota* (**Supplemental Fig. 4**). *Chloroflexota* are abundant and metabolically diverse members of many soil communities but remain comparatively under characterized at the functional level(61). Previous environmental metagenomic studies have identified *Chloroflexota*, *Actinomycetota*, *Bacillota*, *Acidobacteriota*, and *Pseudomonadota* as potential reservoirs of soil ARGs, with novel resistance genes from these diverse lineages also recovered through functional metagenomics(62–64). Thus, the taxonomic breadth observed here is consistent with a broader view of soil resistance as a property distributed across diverse microbial lineages rather than being restricted to recognized antibiotic producers. Importantly, however, our taxonomic assignments represent only the inferred source of short inserts containing candidate sequences and do not establish the host range, mobility, or potential for transfer of these resistance determinants to pathogens.

A major finding of this study is that experimentally validated NTC resistance extends well beyond proteins closely related to previously characterized SATs. SatC (hit 57) and SatD (hit 8) shared maximum amino acid identities of only approximately 31% with characterized SATs, while SatE proteins were also substantially divergent from known references (**Supplemental Fig. 6B**). Previous structural studies identified Glu137 of SatA as a key catalytic residue and Leu136, Ala145, Tyr149, Phe154, and Tyr164 as residues associated with streptothricin binding, and these positions were also used to support the functional assignment of the recently described SatB (21). SatE proteins are predicted to retain an overall SatA-like architecture, whereas SatC and SatD are predicted to show a greater global structural divergence while maintaining a well-superimposed conserved core (**Fig. 8B**). These observations suggest that proteins with substantial sequence divergence can retain structural features associated with streptothricin resistance despite potential differences in their overall structural organization. Additional variation was observed in terminal regions, including a C-terminal extension in the SatE member hit 91 and the previously described N-terminal extension of SatB (**Fig. 8A**). Although the functional significance of these extensions remains unknown, they illustrate additional structural diversity among streptothricin acetyltransferases.

SatD (hit 8) provides a particularly interesting example of this divergence. Most NTC-resistant candidates retained an aspartate corresponding to amino acid residue 160 of SatA, whereas SatD (hit 8) contained an asparagine at this position (**Supplemental Fig. 8A**). Asp160 has previously been implicated as a critical residue in dimerization of *B. anthracis* SatA (59). Specifically, Burckhardt and Escalante-Semerena tested a SatA D160N substitution and found it disrupted proper dimer formation and resulted in protein aggregation, loss of detectable streptothricin binding, and markedly reduced streptothricin-detoxifying activity (MIC reduced from 60 μM down to 2 μM compared to a control strain MIC of 1 μM) (59). Nevertheless, here SatD (hit 8) conferred strong NTC resistance, suggesting that an aspartate at this position may not be universally required across divergent SAT lineages. Structural differences elsewhere in SatD (hit 8) could potentially compensate for this substitution, although this cannot be resolved from the present data without direct oligomerization and biochemical studies. Additional structural features, including the extended C-terminal region of the SatE member hit 91 and the previously described N-terminal extension of SatB(21), may also warrant future investigation (**Fig. 8A**).

The cross-susceptibility analysis revealed further differences among the newly identified SAT lineages. SatC (hit 57) conferred moderate NTC resistance while also significantly increasing the IC_50_ values of amikacin and kanamycin relative to the empty-vector control (**Fig. 6A, and Fig. 7B and 7C**). Notably, however, SatD (hit 8) and SatE (hit 259), which produced substantially stronger NTC resistance phenotypes, did not significantly alter susceptibility to either aminoglycoside. This contrast raises the possibility of a trade-off between strong activity toward streptothricins and broader substrate recognition among divergent acetyltransferases. Such variation is consistent with the diverse substrate specificities of GNAT-family enzymes despite their conserved structural framework(24). Previous studies of aminoglycoside acetyltransferases have further shown that differences in substrate-binding regions can alter antibiotic positioning and catalytic activity(65). However, susceptibility measurements alone cannot establish whether SatC directly acetylates amikacin or kanamycin, or whether these phenotypic differences reflect changes in catalytic specificity. Direct enzymatic assays and product characterization will be required to test this possibility.

Some limitations in our approach should also be considered. Functional metagenomic selection depends on heterologous expression in *E. coli* and may miss genes that are poorly expressed, improperly folded, or dependent on host-specific factors. All NTC-resistance testing performed in our study used ORFs removed from both their local, functionally-selected DNA fragment context and their native genomic context. Consequently, candidates lacking a phenotype in this system cannot necessarily be considered inactive in their native hosts. Furthermore, only a subset of the computationally identified candidates was experimentally examined, leaving additional sequence diversity available for investigation. Finally, the proposed mechanistic relationships are based primarily on sequence conservation and AlphaFold3-predicted structures (**Fig. 8**). Conservation of SatA catalytic and substrate-binding residues is consistent with a shared catalytic mechanism (**Supplementary Fig. 8**) but does not establish that these proteins acetylate the same β-amino group of the streptothricin β-lysine moiety. Direct product characterization, enzyme kinetics, experimental structural studies, and oligomerization measurements will therefore be needed to define the catalytic mechanisms and substrate ranges of SatC, SatD, and SatE.

The SAT diversity identified in this study is consistent with soil microbial communities serving as reservoirs of antibiotic resistance functions that extend beyond currently characterized genes. Streptothricin biosynthesis and self-resistance have been reported in environmental Streptomyces, while the broader soil resistome encompasses diverse and often poorly characterized microbial lineages(66, 67). The identification of SatC, SatD, and SatE expands the recognized diversity of streptothricin resistance-associated acetyltransferases and demonstrates the value of phenotype-based functional metagenomics for discovering divergent resistance determinants. Combined with abundance-weighted sequence analysis and experimental validation, this approach provides a framework for investigating resistance functions that remain poorly captured by conventional sequence annotation. Further characterization of these enzymes will improve our understanding of the environmental streptothricin resistome and may inform the continued development of streptothricins as antibacterial agents.

## Acknowledgments

This work is supported by the Hatch Act of 1887, project award no. 7004080, from the U.S. Department of Agriculture’s National Institute of Food and Agriculture. CRediT contributions: LY, formal analysis, visualization, and writing – original draft. TSC, funding acquisition, project administration, resources, and supervision. LY and TSC, conceptualization, data curation, investigation, and writing – reviewing and editing. See Supplementary figure 9 for a graphical treatment of author contributions. We would like to thank Dr. Carley Reid and Yijun Shi for their insightful comments during the preparation of this manuscript.

## Data availability

Raw PacBio sequencing reads from the NTC selection and sequences for hits are available in NCBI under BioProject PRJNA1550853. Our analysis pipelines and custom code are accessible on github at https://github.com/croftsmicrolab/arg-annotator, and https://github.com/croftsmicrolab/Soil_NTC_paper.

## Notes

### Competing Interest Statement

The authors have declared no competing interest.

